## Supplementary material for "Determinants of species-specific utilization of ACE2 by human and animal coronaviruses": Wang et al Extended data

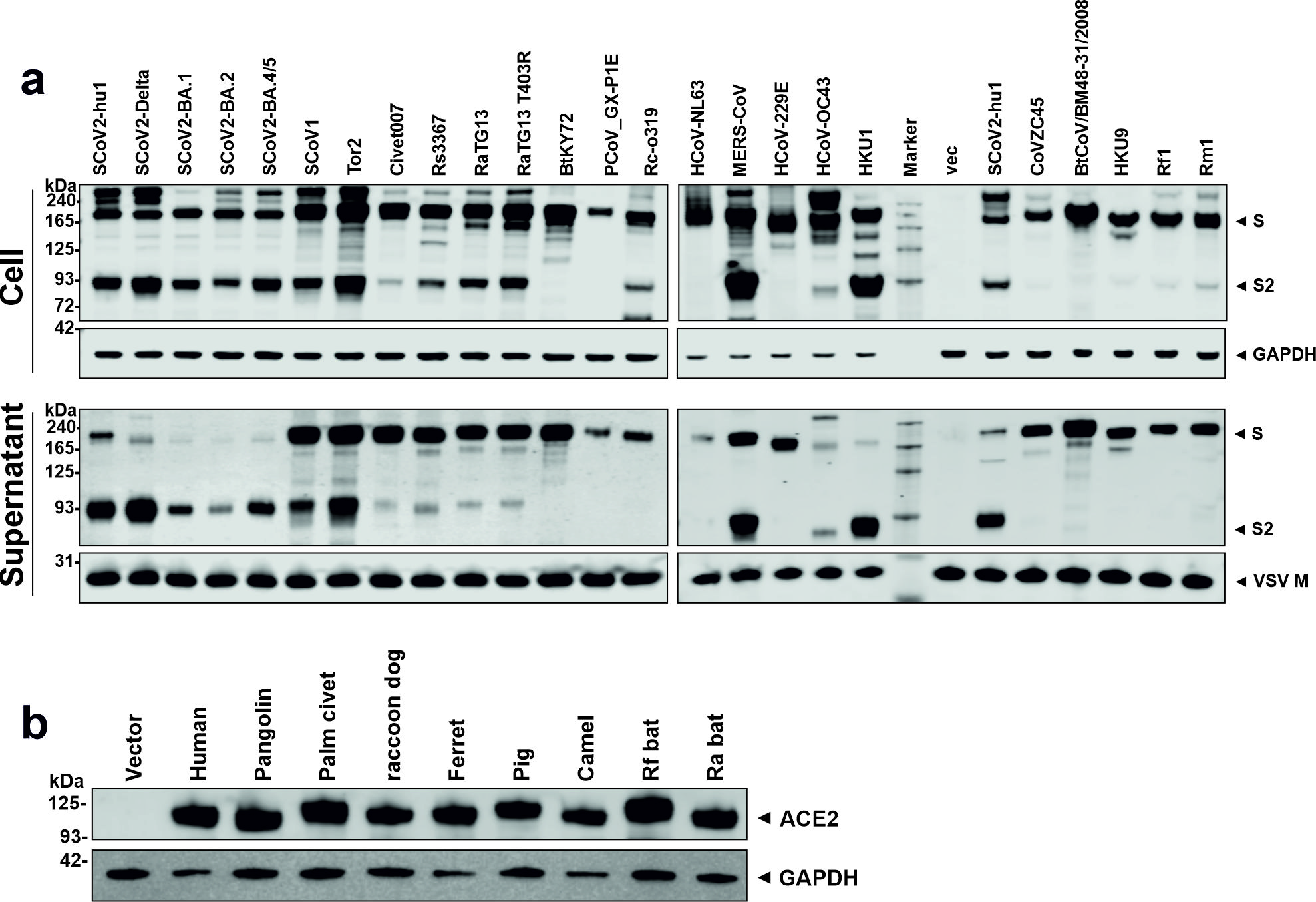

**Figure S1. Expression of S proteins and ACE2 receptors analyzed. a,** Immunoblots of whole cells lysates and supernatants of HEK293T cells transfected with vectors expressing the indicated S proteins. Blots were stained with anti-strep tag, anti-GAPDH, and anti-VSV-M. **b**, Western blot analyses of extracts of HEK293T cells transfected with expression constructs for the indicated ACE2 orthologs or an empty control vector.

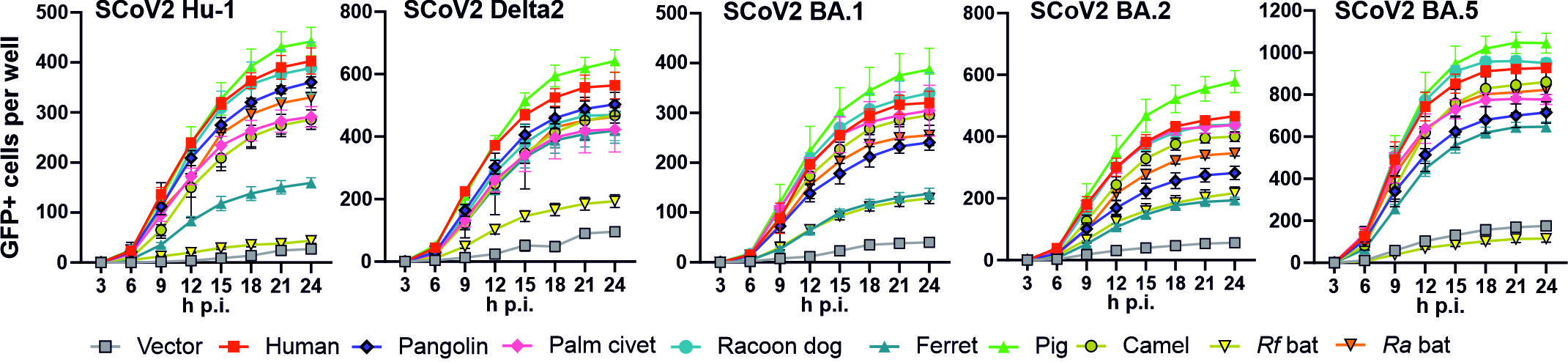
 **Figure S2. Utilization of different ACE2 orthologs by the S proteins of SARS-CoV-2 variants.** Infection kinetics of HEK293T cells expressing ACE2 receptors from the indicated species after exposure to VSVpp containing the S proteins of SARS-CoV-2 variants. Infected GFP+ cells were automatically quantified over a period of 24 h.

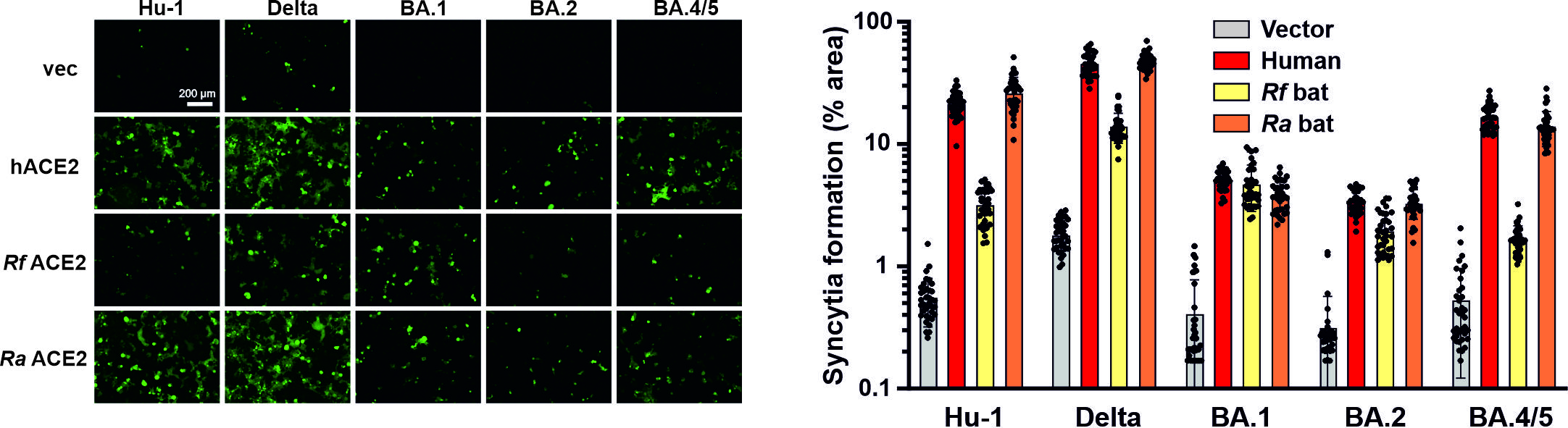

**Figure S3. BA.5 S does not allow *Rf* ACE2 dependent cell fusion**. Left, Exemplary fluorescence microscopy images of HEK293T cells expressing SARS-CoV-2 S protein and the indicated ACE2 orthologs (left). Scale bar, 200 μm. The right panel shows the automatic quantification of syncytia formation by calculating the GFP positive areas. Bars represent the mean of three independent experiments (±SEM).

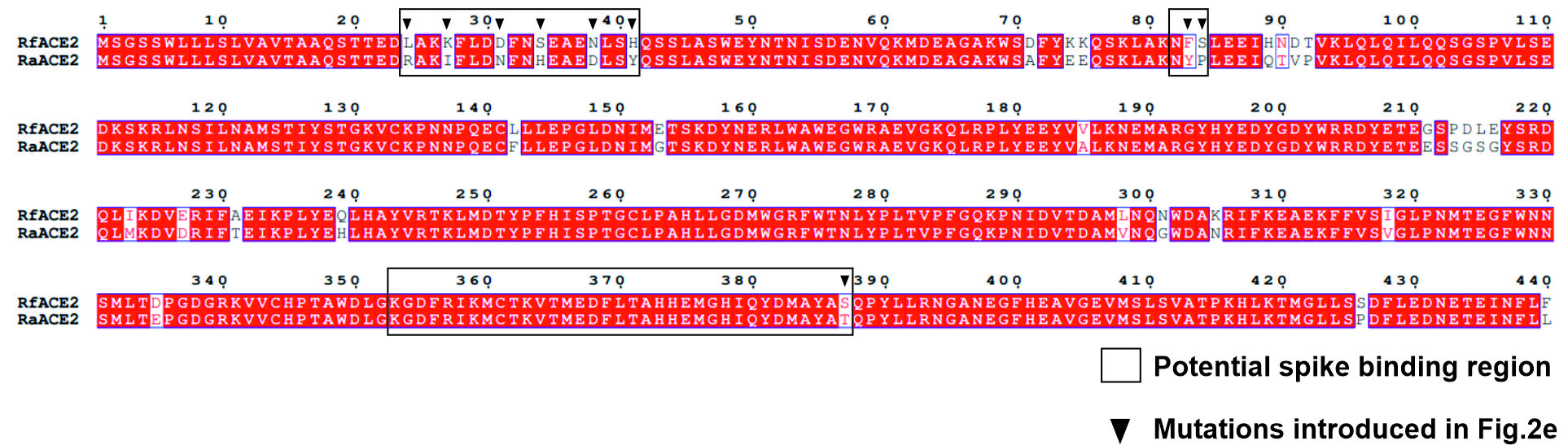

**Figure S4. Mutations introduced into *Rf* ACE2.** Conserved areas are indicated in red. Number give amino acid positions in the *Rf* and *Ra* ACE2 sequence.

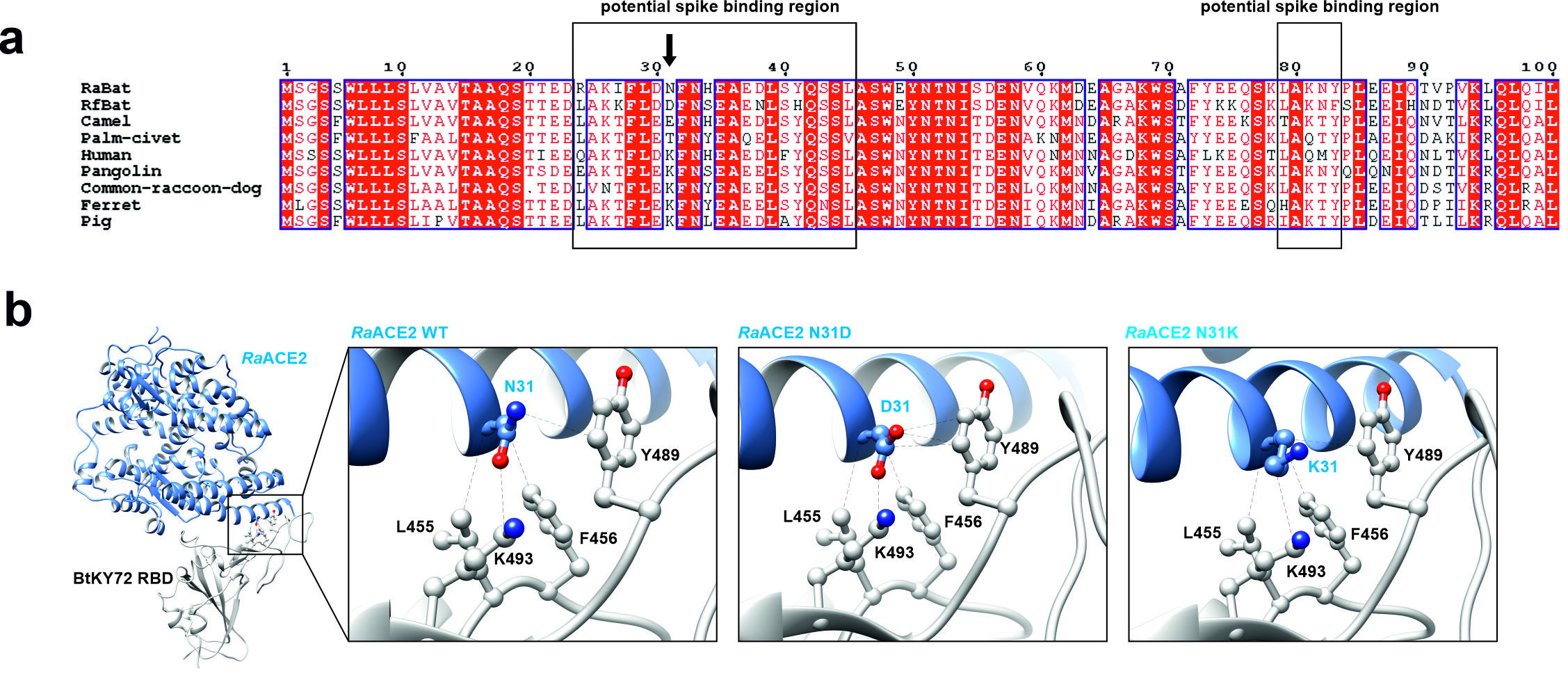
 **Figure S5. Potential determinants of species-Specificity of ACE2 receptor function. a,** Alignment of ACE2 sequences analyzed. **b.** Schematic diagram of BtKY72 RBD with the *Ra* ACE2. Potential van der Waals interactions of BtKY72 RBD with the *Ra* ACE2 wide-type, N31D or N31K are indicated by dash back lines.

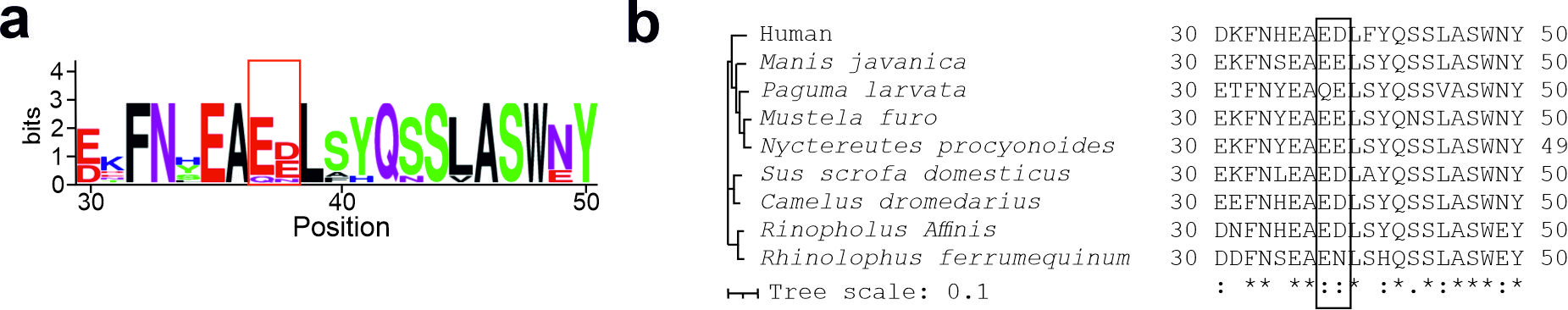

**Figure S6. Conservation of acid amino acids at positions 37 and 38 of the ACE2 orthologs analyzed.** **a,** Sequence logo of the alignment of ACE2 sequences between sequence positions 30 and 50. Positions 37 and 38 are highlighted by a red box. **b,** Alignment of amino acid residues 30 to 50 of ACE2 sequences analyzed.

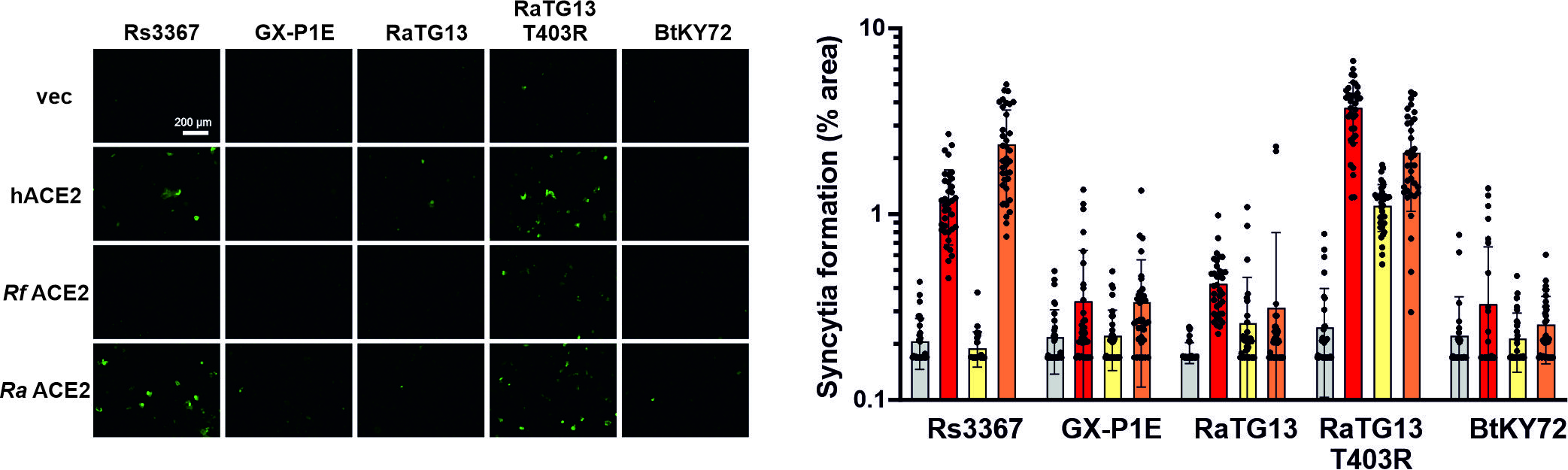

**Figure S7. Species-specific ability of bat CoV S proteins for ACE2-mediated cell-to-cell fusion.** Fluorescence microscopy images of HEK293T cells expressing S proteins of bat CoVs and the indicated ACE2 orthologs (left). Scale bar, 200 μm. The right panel shows shows the automatic quantification of syncytia formation by calculating the GFP positive areas. Bars represent the mean of three independent experiments (±SEM). Statistical significance was tested by two-tailed Student’s t test with Welch’s correction. *p < 0.05; **p < 0.01; ***p < 0.001.

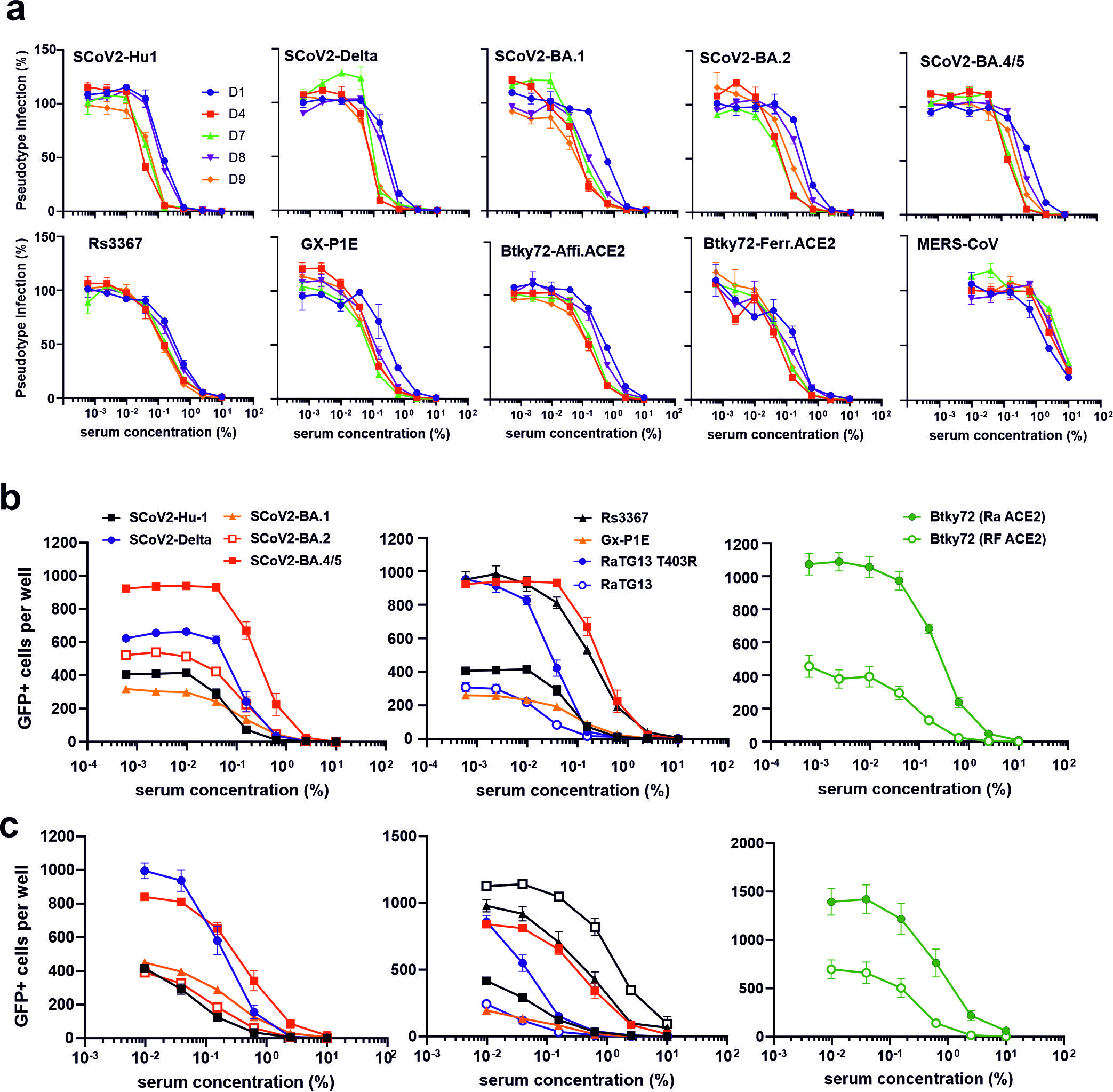

**Figure S8. Neutralization of VSVpp infection mediated by S proteins of human and bat CoVs. a,** Neutralization of VSVΔG-GFP pseudotyped with the indicated S proteins by sera from five AZ/2xBNT162b2 vaccinated individuals. Lines represent the mean of three biological replicates. **b, c**, Infection of HEK293T cells expressing human or bat (right panels) ACE2 receptors by VSVpp carrying the S proteins of the indicated human and bat CoVs in the presence sera obtained from five AZ/BNT/BNT (b) or five 3xBNT (c) vaccinated individuals. Shown are total numbers of VSVpp infected (GFP+) cells in the presence of the indicated concentrations of sera. Shown are mean values obtained for the five sera, each tested in two technical replicates.

**Table S1. Spike sequences analyzed.**

| **Spike Name** | **Origin Lat. Name** | **Seq. identity to SCoV2 S Hu1 (%)** | **Accession Nr.** |
| --- | --- | --- | --- |
| SARS-CoV-2 Hu1 | human | 100 | BCN86353.1 |
| SARS-CoV-2 Delta | human | 99,21 | QTW89558.1 |
| SARS-CoV-2 Omicron BA.1 | human | 96,94 | UFO69279.1 |
| SARS-CoV-2 Omicron BA.2 | human | 97,56 | UHU97100.1 |
| SARS-CoV-2 Omicron BA.4/5 | human | 97,33 | UPN16705.1 |
| RATG13 | *Rh. affinis* | 97,41 | QHR63300.2 |
| GX/PIE | *M. javanica* | 92,73 | QIA48623.1 |
| CoVZC45 | *Rh. sinicus* | 80,69 | AVP78031.1 |
| SARS-CoV-1 ShanghaiQXC2 | human | 78,88 | AAR86775.1 |
| Rs3367 | *Rh. sinicus* | 77,07 | AGZ48818.1 |
| Rc-o319 | *Rh. cornutus* | 76,48 | BCG66627.1 |
| SARS-CoV-1 TOR2 | human | 76,04 | P59594.1 |
| Civet007 | *P. larvata* | 75,8 | AAU04646.1 |
| Rm1 | *Rh. macrotis* | 74,75 | ABD75332.1 |
| Rf1 | *Rh. ferrumequinum* | 74,00 | ABD75323.1 |
| BtKY72 | *Rh. sp.* | 72,03 | APO40579.1 |
| BM48-31/2008 | *Rh. blasii* | 62,99 | YP_003858584.1 |
| OC43 | human | 37,93 | AVR40344.1 |
| HKU1 | human | 35,95 | YP_173238.1 |
| MERS | human | 34,82 | QBM11748.1 |
| HKU9 | *Rousettus sp.* | 33,33 | AVP25406.1 |
| NL63 | human | 31,27 | YP_003767.1 |
| 229E | human | 31,11 | NP_073551.1 |

**Table S2. ACE2 sequences analyzed.**

| **ACE2 Species Name** | **Lat. Name** | **Seq. identity to human ACE2 (%)** | **Acession Nr.** |
| --- | --- | --- | --- |
| Human | *Homo Sapiens* | 100 | BAB40370.1 |
| Pangolin | *Manis javanica* | 84,63 | XP_017505752.1 |
| Palm Civet | *Paguma larvata* | 83,50 | Q56NL1.1 |
| Raccoon Dog | *Nyctereutes procyonoides* | 83,75 | ABW16956.1 |
| Ferret | *Mustela putorius furo* | 82,49 | NP_001297119.1 |
| Pig | *Sus scrofa domesticus* | 81,74 | XP_020935033.1 |
| Camel | *Camelus ferrus* | 83,12 | XP_006194263.1 |
| *R. ferrumequinum* | *Rh. Ferrumequinum* | 81,23 | XP_032963186.1 |
| *R. affinis* | *Rh. affinis* | 80,60 | QMQ39229.1 |
